# Multi-modality Graph Representation Learning for Malignant Cell Identification from scRNA-seq using DeepMalignant

**DOI:** 10.64898/2026.06.29.734828

**Authors:** Pankaj Bhattarai, Weiman Yuan, Hongmei Chi, Xin Maizie Zhou, Xian Mallory

## Abstract

Distinguishing malignant from normal cells in single-cell RNA sequencing data remains a critical yet challenging task in cancer genomics. Existing methods often suffer from poor precision, limited generalizability across cancer types, and reduced robustness across different sequencing platforms. We developed DeepMalignant, an unsupervised multimodal graph attention autoencoder for malignant cell identification that jointly integrates gene expression and copy number alteration (CNA) information. We applied DeepMalignant to five datasets covering 26 samples and four cancer types (breast, colorectal, pancreatic, and ovarian cancers), generated by three platforms (10x Genomics, inDrop, and Drop-seq) for benchmarking and compared it with existing state-of-the-art methods including scMalignantFinder, PreCanCell, CopyKAT, ikarus, and Cancer-Finder. DeepMalignant achieved the best overall balance of precision and recall and consistently outperformed the existing methods that used either gene expression or CNA in F1 scores. Ablation studies showed that both CNA-based edge weighting and graph attention aggregation contribute independently to performance, and attribution analysis further indicated that the learned embeddings capture biologically meaningful malignant programs. We further applied DeepMalignant to two ductal carcinoma in situ (DCIS) samples, DCIS2 and DCIS1, that have matched spatial transcriptomics and scRNA-seq data. DeepMalignant identified tumor-enriched regions that were highly consistent with the matched histological image. The downstream cellcell communications analysis revealed that fibroblast-derived C3 and MIF both directed signaling more toward normal epithelial cells than tumor epithelial cells, demonstrating that accurate tumor-normal cell classification by DeepMalignant enables biologically meaningful interrogation of the tumor microenvironment and revealing how stromal cells differentially communicate with malignant versus normal epithelial populations.

---

Cancer develops through the progressive accumulation of molecular alterations, yet tumors are not uniform cell populations. Instead, each tumor contains a heterogeneous mixture of malignant cells together with fibroblasts, endothelial cells, and diverse immune populations [1, 2]. Single-cell RNA sequencing (scRNA-seq) has made this heterogeneity directly measurable. However, cells sequenced by scRNA-seq do not reveal cell identity such as malignant or normal cells naturally [1, 3, 4]. Accurately and reproducibly identifying malignant and normal cells is fundamental in the downstream analysis to prevent errors in this step from propagating into differential expression analysis, subclone analysis, microenvironment profiling, and any attempt to interpret tumor evolution or cell-cell interactions [1, 2, 4].

Current methods for malignant cell annotation from scRNA-seq fall into two broad groups. One group infers copy number alteration (CNA) patterns from transcriptomic data and uses aneuploidy as evidence of malignancy. CopyKAT is a leading example: it estimates broad CNA profiles from scRNA-seq and separates aneuploid from diploid cells while also recovering clonal structure [1]. This strategy is effective in many solid tumors, but CNA-based inference alone is insufficient. It can degrade in near-diploid tumors and samples with few normal reference cells, and when CNA signal is not pronounced, as is the case in tumors with low somatic copy-number burden [1, 5–7]. Recent benchmarking studies further demonstrate that the performance of CNA inference methods is highly sensitive to sequencing platform, read depth, tumor purity, dropout rate, and the choice of reference cells — with broad chromosomal-scale alterations recovered far more reliably than focal copy-number events [5–7]. In tumors with low tumor purity and modest or near-diploid copy-number profiles, such as pancreatic ductal adenocarcinoma (PDAC), different CNA-based callers can disagree substantially, often yielding inconsistent malignancy labels and an elevated rate of false positives [8].

A second family of methods uses gene expression directly. Ikarus builds cross-dataset malignant signatures and combines logistic regression with label propagation to classify cells [3]. PreCanCell uses an ensemble of classifiers trained on malignant and non-malignant marker genes shared across several cancer types [9]. Cancer-Finder uses a domain generalization framework to improve transfer across tissues and extends the same idea to spatial transcriptomic data [10]. ScMalignantFinder uses curated cancer signatures and a large calibrated pan-cancer training set to capture broader malignant transcriptional diversity [4]. These methods are often highly sensitive, but limited by training data composition and shifts in malignant transcriptional programs across tissues, datasets, and disease states [3, 4, 9, 10]. The limitation is often manifested by low specificity in calling malignant cells especially when tumor and normal populations have overlapping expression programs. As reported in the scMalignantFinder benchmarking [4] on the scRNA-seq datasets from [11, 12], scMalignantFinder achieved the highest specificity of 0.786 among compared methods, compared with 0.642 for ikarus, 0.503 for PreCanCell, 0.397 for CopyKAT, and 0.022 for Cancer-Finder. Yet even the highest specificity of 0.786 falls short of the specificity required for reliable downstream analysis.

To overcome these limitations, research has shifted toward deep learning architectures capable of capturing high-order biological dependencies. Recently, the Transcriptome Graph Transformer (TGT) demonstrated that unsupervised pre-training on heterogeneous gene-pathway graphs can significantly enhance the representation of transcriptomic data [13]. Furthermore, Attention-Based Hierarchical Graph Autoencoders have proven effective at resolving single-cell resistance dynamics by integrating biological hierarchies with attention mechanisms [14], emphasizing the power of graph neural networks in modeling complex cellular trajectories. However, a gap remains in explicitly leveraging both genomic structural signals, i.e., CNAs, and transcriptomic signatures within a unified graph framework to achieve robust cell-type identification.

We developed DeepMalignant, a multimodal deep learning framework that integrates both gene expression and CNAs to achieve more accurate discrimination of malignant from non-malignant cells. DeepMalignant is formulated as a graph neural network in which nodes represent individual cells, node features encode RNA-derived cancer signature profiles, and edge weights capture CNA-derived pairwise cell similarity. The model then learns latent cell embeddings through a graph attention autoencoder, enabling it to selectively attend to the most informative cellular relationships during representation learning. The graph attention network naturally combines the strengths of both signal types: gene expression captures malignant cell state, while CNA similarity captures shared genomic architecture that is often more stable across cells from the same tumor clone. Critically, CNA-informed edge weights enable cells with similar chromosomal profiles to mutually reinforce one another during representation learning, mitigating the elevated false positive rates inherent to gene-expression-only methods while simultaneously compensating for the instability of CNA-only inference such as in low-purity samples and tumors with modest copy-number alterations.

We benchmarked DeepMalignant against five state-of-the-art methods — scMalignantFinder, PreCanCell, CopyKAT, ikarus, and CancerFinder — across five datasets encompassing 26 samples, four cancer types (breast, colorectal, pancreatic, and ovarian), and three sequencing platforms (10x Genomics, inDrop, and Drop-seq). Across this heterogeneous collection, DeepMalignant consistently achieved among the highest F1 scores, reflecting a superior balance of precision and recall, with particularly pronounced advantages in samples where RNA-only methods overpredicted malignant cells or where CopyKAT exhibited marked instability across samples.

## Results

### Method overview

**Figure 1** provides an overview of the DeepMalignant workflow and downstream analyses. DeepMalignant takes single-cell RNA-seq data as input, including a gene expression matrix and an inferred copy number alteration (CNA) matrix (**Figure 1A**). The gene expression matrix is used to define RNA-derived node features based on curated cancer-associated signature genes, while the CNA matrix is used to construct CNA-informed cell–cell relationships. Specifically, pairwise CNA similarity is used to define weighted edges between cells, producing a graph in which each node represents a cell and each edge reflects genomic similarity between cells (**Figure 1B**). This multimodal graph is then passed into a graph attention autoencoder, where the encoder learns low-dimensional cell embeddings through CNA-weighted message passing and the decoder reconstructs the input features to stabilize representation learning (**Figure 1C**). The learned embeddings are subsequently clustered in the low-dimensional latent space using the Leiden community detection algorithm [15] (**Figure 1D**). To assign biological labels to these clusters, DeepMalignant computes cluster-level CNA burden scores and fits a two-component Gaussian mixture model, where clusters with higher CNA burden are classified as malignant and the remaining clusters are classified as normal (**Figure 1E–F**). Beyond malignant cell identification, the learned model also supports downstream interpretation and applications. We quantify the relative contribution of CNA and cancer-related transcriptional pathways using perturbation-based importance scores and we further apply DeepMalignant to spatial transcriptomics data to identify tumor-enriched regions and normal-spot cell-type compositions, and characterize the cell-cell communications (**Figure 1G-I**).

**Figure 1:**
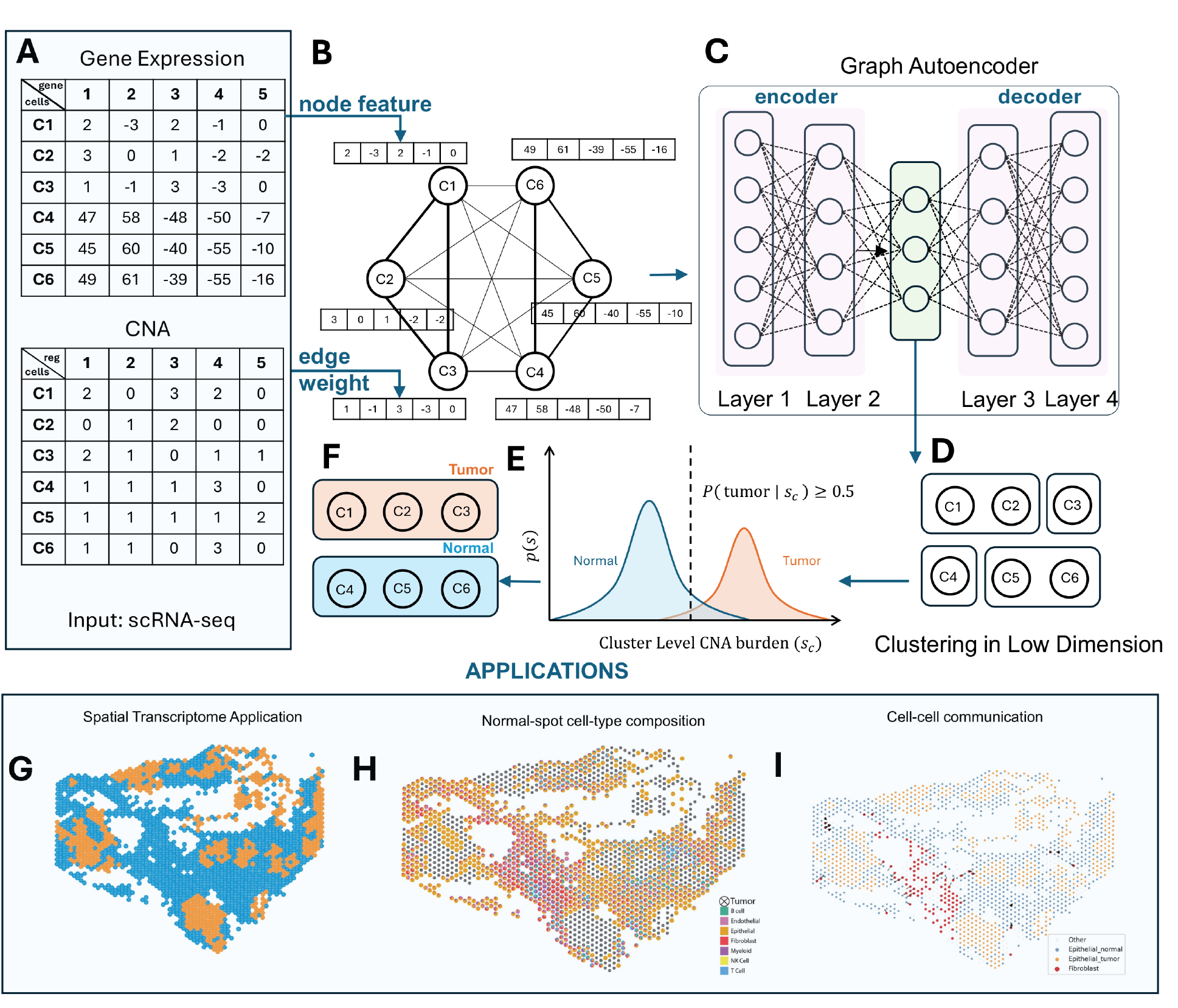
Overview of the proposed multi-omic graph attention framework. (A) Gene expression matrix provides RNA-based node features. (B) CNA matrix is used to construct weighted cell–cell edges via cosine similarity. (C) A graph attention autoencoder learns low-dimensional embeddings using CNA-weighted message passing. (D) Latent embeddings are clustered to identify malignant and normal cell populations. (E) Malignant-normal classification via a two-component Gaussian mixture model fitted on cluster level CNA burden. (F) Final classification into malignant and normal cells. (G) Spatial transcriptome application of DeepMalignant, showing the spatial distribution of predicted tumor and normal spots. (H) Cell-type composition of spots predicted as normal, illustrating the inferred contribution of major cell populations within the normal spatial compartment. (I) Cell–cell communication analysis in the spatial transcriptome data, highlighting interactions between fibroblasts and epithelial normal or tumor spots.

### Overview of benchmarking

We thoroughly evaluated the precision, recall and accuracy (F1 score) of DeepMalignant by applying it to multiple scRNA-seq data, spanning four cancer types (breast, ovarian, colorectal and pancreatic), three sequencing technologies (10x Genomics, Drop-seq, and inDrop), and a total of 26 samples (details of the datasets seen in **SI Appendix Table 1**). These datasets were obtained from the Curated Cancer Cell Atlas, which provided the ground truth labeling of the malignant cells. We benchmarked DeepMalignant with five state-of-the-art methods, including scMalignantFinder, PreCanCell, ikarus, Cancer-Finder, and CopyKAT. Detailed results for each sample and each method are reported in **Dataset S1**.

**Table 1:**
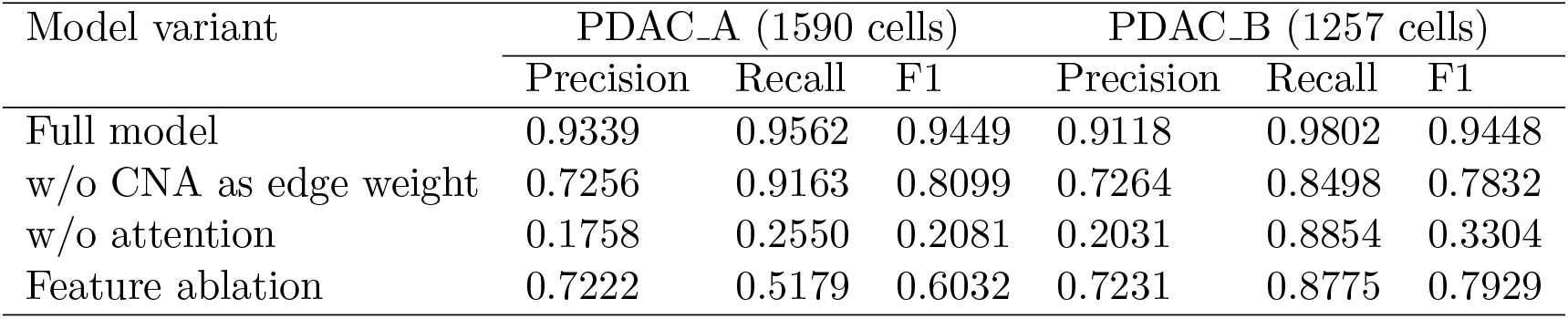
Ablation analysis of DeepMalignant on the Moncada et al. dataset. The full model was compared with three ablated variants: constant edge weights, an expression-only baseline without graph attention, and a feature-ablation model in which the union of the top 20 genes most strongly correlated with each latent dimension was removed from the input feature set. Precision, recall, and F1 score are reported for each sample.

**Figure 2A** presents the recall, precision, and F1 scores across all five benchmark datasets, providing an overview of the six methods’ performance, including DeepMalignant. Among the six methods, ikarus performs the worst, with median recall and F1 scores of 0. ScMalignantFinder exhibits the lowest precision overall; its characteristic pattern of high recall coupled with low precision has been reported previously [4]. PreCanCell shows a similar performance profile to scMalignantFinder, though with somewhat higher precision. CopyKAT, CancerFinder, and DeepMalignant are substantially more competitive than ikarus, scMalignantFinder, and PreCanCell. Although CopyKAT achieves median precision (0.9651) and F1 (0.9528) scores slightly higher than DeepMalignant’s (0.9519, 0.9459), its median recall (0.9819) is lower (0.9909), and it yields zero F1 scores for several samples, likely attributable to an incorrect diploid baseline selection. CancerFinder achieves the highest recall of all methods; however, its median precision (0.8956) falls below that of DeepMalignant (0.9519), and its median F1 score (0.9437), while close, remains lower than DeepMalignant’s (0.9459). Overall, DeepMalignant achieved the second highest median F1 scores while maintaining the best balance between recall and precision and robustness. The following sections examine each method’s performance on individual datasets, organized by cancer type and sequencing platform, to provide a more granular understanding of each method’s strengths and limitations. **SI Appendix: details of running existing methods** listed the benchmarking implementation details for each method.

**Figure 2:**
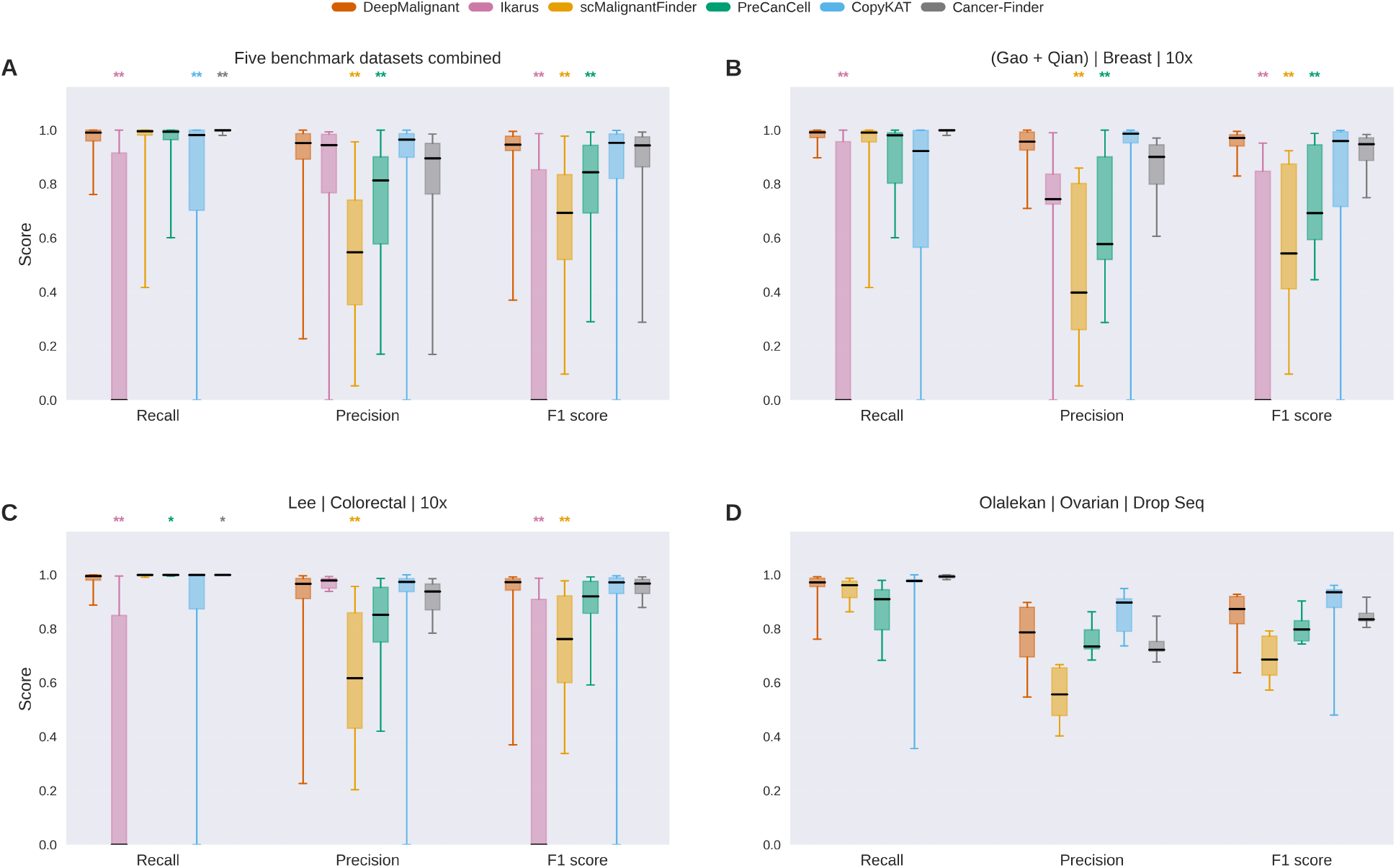
Cross-dataset benchmarking of DeepMalignant for malignant cell identification in single-cell transcriptomic data. Boxplots summarize per-sample recall, precision, and F1 score for DeepMalignant and existing state-of-the-art methods. **A.** Results pooled across five benchmark datasets: Moncada et al. (pancreatic cancer, inDrop), Gao et al. and Qian et al. (breast cancer, 10x), Lee et al. (colorectal cancer, 10x), and Olalekan et al. (ovarian cancer, Drop-seq). **B**. Combined breast cancer 10x datasets from Gao and Qian et al. **C**. Lee et al. colorectal cancer 10x dataset. **D**. Olalekan et al. ovarian cancer Drop-seq dataset. Asterisks denote paired Student’s *t*-test significance for comparisons between DeepMalignant and competing methods (*, *p <* 0.05; **, *p <* 0.01).

### DeepMalignant achieved accurate and robust malignant cell identification in breast and colorectal cancer datasets sequenced by 10x Genomics

We first applied DeepMalignant to two breast cancer scRNA-seq data generated by 10x Genomics: Gao et al. [1] and Qian et al. [16], comprising in total 11 breast cancer samples and 20,673 cells. The Gao et al. dataset includes one ductal carcinoma in situ sample (DCIS1) and three triple-negative breast cancer samples (TNBC1–TNBC3), whereas the Qian et al. dataset includes seven additional breast cancer samples. For both datasets, ground-truth malignant and normal cell labels were obtained from the benchmark annotations, and gene expression matrices were preprocessed using standard filtering and normalization procedures before graph construction.

For these two breast cancer datasets, we evaluated DeepMalignant and the five state-of-the-art methods using per-sample precision, recall, and F1 score for malignant cells, as shown in **Figure 2B**. DeepMalignant consistently achieved the highest F1 score (median F1: 0.9713). CopyKAT and CancerFinder ranked second and third (median F1: 0.9600 and 0.9480, respectively), while PreCanCell, scMalignantFinder, and Ikarus all performed substantially worse (median F1 *<* 0.7 for all three). On this dataset, CopyKAT’s F1 score exhibited considerably higher variance (0.1097) than DeepMalignant’s (0.0022), and one sample (Qian et al., sample 42) yielded a zero F1 score, indicative of an incorrect diploid baseline selection by CopyKAT. CancerFinder achieved the highest recall of all methods (median: 0.9997), marginally exceeding DeepMalignant (median: 0.9923); by contrast, DeepMalignant’s precision (median: 0.9575) was substantially higher than that of CancerFinder (median: 0.9011). Although CopyKAT attained the highest median precision overall, it produced a zero precision score for sample 42 in Qian et al., undermining its robustness. Overall, on the Gao et al. and Qian et al. breast cancer datasets sequenced with 10x, DeepMalignant achieved the highest F1 score and the best balance between recall and precision.

We then evaluated DeepMalignant on the Lee et al. [17] colorectal cancer scRNA-seq dataset generated by the 10x Genomics technology. This dataset comprises 14 colorectal cancer samples and 20,977 cells. DeepMalignant showed favorable performance over most state-of-the-art methods for this dataset (**Figures 2C**). Particularly, DeepMalignant achieved the second highest median F1 score (0.973) among the six methods, only next to CopyKAT (0.9818). Although DeepMalignant suffered from the low precision for one colorectal sample (precision = 0.2268, F1 score = 0.3697) in this dataset, all other samples’ precision and F1 score maintained to be high (precision ≥ 0.7394, F1 ≥ 0.8502). CopyKAT suffered from low F1 score for two samples (0 and 0.3553), showing that it is not as robust as DeepMalignant in this colorectal cancer dataset. We further investigated the sample that CopyKAT’s F1 was zero (sample SMC18), and found that CopyKAT reported that they encountered “low confidence in classification”, showing that it could not confidently establish a diploid baseline. When CopyKAT selected the wrong baseline cluster, its aneuploid and diploid labels were inverted, leading to near zero F1 score for the sample. This showed that CopyKAT was heavily constrained by its selection of the normal baseline, which was subject to the number of normal cells in the sample and the CNA profiles of the normal cells.

Overall, DeepMalignant obtained the highest accuracy and consistency across the 10x Genomics breast cancer and colorectal datasets. Cancer-Finder and CopyKAT were the next most competitive methods. However, CopyKAT suffered from poor recall, precision, and F1 score for some outlier samples in both Qian’s and Lee’s datasets. It also heavily relied on the correct selection of the normal baseline. Cancer-Finder, on the other hand, had lower median precision and F1 score than DeepMalignant in all three datasets.

### DeepMalignant was robust to ovarian cancer samples sequenced by Drop-seq

We further evaluated DeepMalignant on a metastatic ovarian cancer dataset sequenced by Drop-seq from Olalekan et al. [18]. This dataset contains six samples comprising a total of 6,018 cells. We applied the six methods (DeepMalignant, ikarus, scMalignantFinder, PreCanCell, CopyKAT and Cancer-Finder), of which ikarus could not be successfully run on this dataset using the default pipeline recommended by the authors and was therefore excluded from the quantitative comparison.

Of all five methods, DeepMalignant demonstrated robust, strong and balanced performance in between recall and precision in this ovarian cancer dataset (**Figures 2D**). Its median F1 score (0.873) was the second highest after CopyKAT (0.936). Nevertheless, CopyKAT suffered poor recall on outlier samples, and its lowest F1 score was lower than 0.5, possibly because it relied solely on the copy number signals. In contrast, DeepMalignant demonstrated a good balance between recall and precision, thanks to it incorporating both the gene expression and copy number signals. Like those datasets in 10x Genomics, scMalignantFinder and PreCanCell showed less favorable results than DeepMalignant in recall, precision and F1 score in this dataset. Cancer-Finder demonstrated high recall overall, but its median F1 score (0.836) is lower than that of DeepMalignant (0.8731).

In all, DeepMalignant demonstrated its ability to be applied to Drop-seq dataset on the metastatic Ovarian cancer dataset, featuring its balanced recall and precision, robustness and high F1 score.

### DeepMalignant uniquely maintained robust malignant cell identification in a pancreatic cancer dataset sequenced by inDrop sequencing

To further evaluate DeepMalignant on a distinct tumor type and sequencing platform, we applied DeepMalignant and other five state-of-the-art methods to a pancreatic ductal adenocarcinoma (PDAC) single-cell dataset generated using inDrop technology from Moncada et al. [19]. This dataset contains two pancreatic cancer samples, PDAC_A and PDAC_B, comprising 1,590 and 1,257 cells, respectively.

Since there were only two samples, we displayed the recall and precision using a radar plot (**Figure 3A**). In this dataset, DeepMalignant uniquely showed strong and highly balanced performance for both PDAC samples, achieving nearly identical F1 scores of 0.9449 in PDAC_A and 0.9448 in PDAC_B. In contrast, CopyKAT failed in PDAC_B on both recall and precision and its F1 score dropped to 0.0073. Other methods such as PreCanCell, Cancer-Finder and scMalignant all displayed high recall for both PDAC_A and PDAC_B, but low precision, showing on the radar plot as more a diamond shape than a square. On this dataset, ikarus could not be successfully run using the default pipeline recommended by the authors and was therefore excluded from the comparative evaluation.

**Figure 3:**
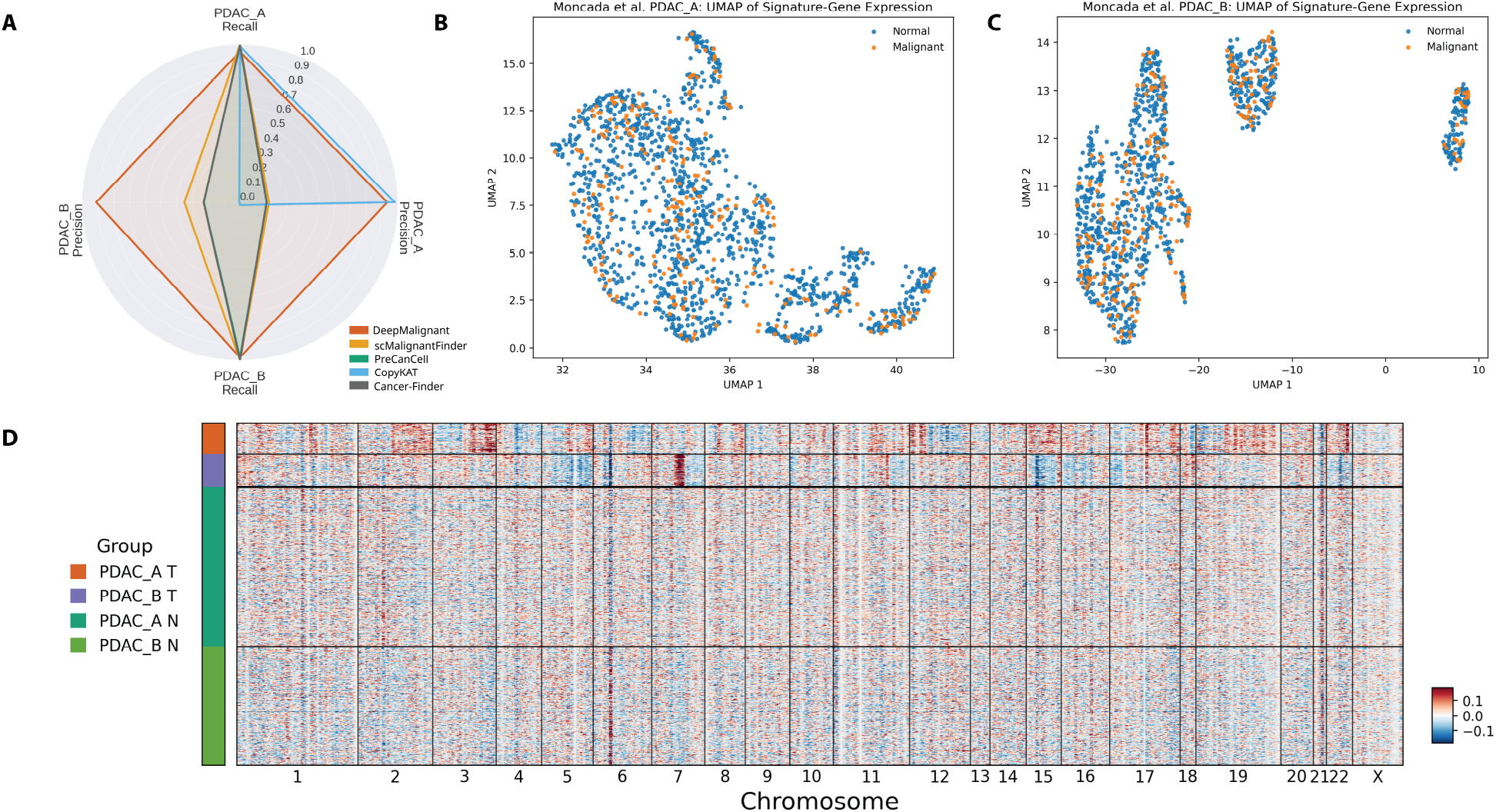
Performance and transcriptomic characterization of tumor cell identification in the Moncada et al. pancreatic cancer dataset. (A) Radar plot comparing recall and precision across PDAC_A and PDAC_B samples for DeepMalignant and baseline methods. (B–C) UMAP visualization of signature-gene expression space for PDAC_A and PDAC_B, respectively, colored by ground truth labels, showing limited separation between malignant and normal cells. (D) Genome-ordered heatmap of normalized expression across all chromosomes for individual cells. Cells are grouped by samples and predicted classes (red: PDAC_A tumor; blue: PDAC_B tumor; dark green: PDAC_A normal, and light green: PDAC_B normal). Genes were first ordered according to their genomic coordinates and expression values were normalized using counts per 10,000 followed by log-transformation. Expression values were then centered per gene across all cells and smoothed using a sliding window of 50 consecutive genome-ordered genes.

To elucidate the failure of state-of-the-art methods on the two pancreatic cancer samples, we visualized the UMAP projections of the input signature-gene set for both PDAC_A (**Figure 3B**) and PDAC_B (**Figure 3C**). In both cases, the transcriptomic profiles of malignant cells substantially overlapped with those of normal cells, illustrating the inherent difficulty of relying on transcriptomic features alone for malignant cell identification in this dataset. By contrast, the Gao et al. breast cancer samples (DCIS1, TNBC1–3) exhibited markedly clearer separation between malignant and normal cells, with well-defined cluster boundaries observed consistently across all four samples (**SI Appendix Fig. S1**). Collectively, these results demonstrated that transcriptomic differences between malignant and normal cells were considerably more ambiguous in the Moncada pancreatic dataset than in the breast cancer cohort, reinforcing the necessity of incorporating complementary modalities beyond gene expression for robust malignant cell identification.

To investigate the failure of CopyKAT on PDAC_B, we examined the normalized expression counts for both samples (**Figure 3D**). Analysis of PDAC_B revealed sparse copy number variation across the genome, characterized by a single amplification on chromosome 7 and isolated deletions on chromosomes 5, 15, 16, 17, and 22. We attributed the poor performance of existing methods to the concomitant weakness of both transcriptomic and copy number variation signals in this sample, which individually proved insufficient for reliable malignant cell identification. These findings underscored a key advantage of DeepMalignant: by jointly leveraging both modalities, it maintained robust discrimination between malignant and normal cells even when either signal alone was uninformative.

### DeepMalignant correctly identified malignant and normal spots in two spatial transcriptomics DCIS samples

We further evaluated DeepMalignant’s capacity of generalizing beyond scRNA-seq to spatial transcriptomics data by applying it to two ductal carcinoma in situ (DCIS) samples, DCIS1 and DCIS2, from the study of Wei et al. [20], which were profiled using the 10x Genomics Visium platform that included the spatial coordinates for each spot.

We presented the spatial distributions of malignant (orange) and normal (blue) predictions for the DCIS2 sample across DeepMalignant (**Figure 4A**), DeepMalignant with constant edge weights (**Figure 4B**), and four existing methods (**Figure 4C–F**). The constant-edge-weight variant of DeepMalignant was included to examine the contribution of copy number profiles to the overall model performance in DeepMalignant. For reference, the corresponding histopathology image with expert-annotated tumor regions, adapted from [20], is shown in **Figure 4H**, corresponding to the subregion delineated by the red box in **Figure 4A**. As described in [20], this subregion encompasses eleven annotated tumor regions (T1–T11). The boxed region from **Figure 4A** was then enlarged to match the size of the histopathology image in **Figure 4H**, resulting in **Figure 4G**. On the same figure, we also manually grouped the malignant spots based on their positions while comparing with the expert annotation from the histopathology image in **Figure 4H**, which showed that DeepMalignant’s predictions of malignant spots for DCIS2 had near-perfect concordance with the histopathology-derived annotations. Examination of **Figure 4B–F** revealed that competing methods performed substantially worse. Removing copy number profiles from edge weights in the constant-edge-weight variant disrupted spatial coherence among malignant spots, yielding fragmented prediction regions. CopyKAT (**Figure 4C**) and Cancer-Finder (**Figure 4E**) exhibited a systematic tendency toward over-calling malignant spots, erroneously merging T5 with T6 and T7 through T9 into contiguous regions. This over-calling of malignant spot was further exacerbated in scMalignantFinder and PreCanCell, with scMalignantFinder classifying nearly all spots as malignant. Taken together, DeepMalignant achieved the highest spatial accuracy in malignant spot identification on the DCIS2 10x Visium dataset.

**Figure 4:**
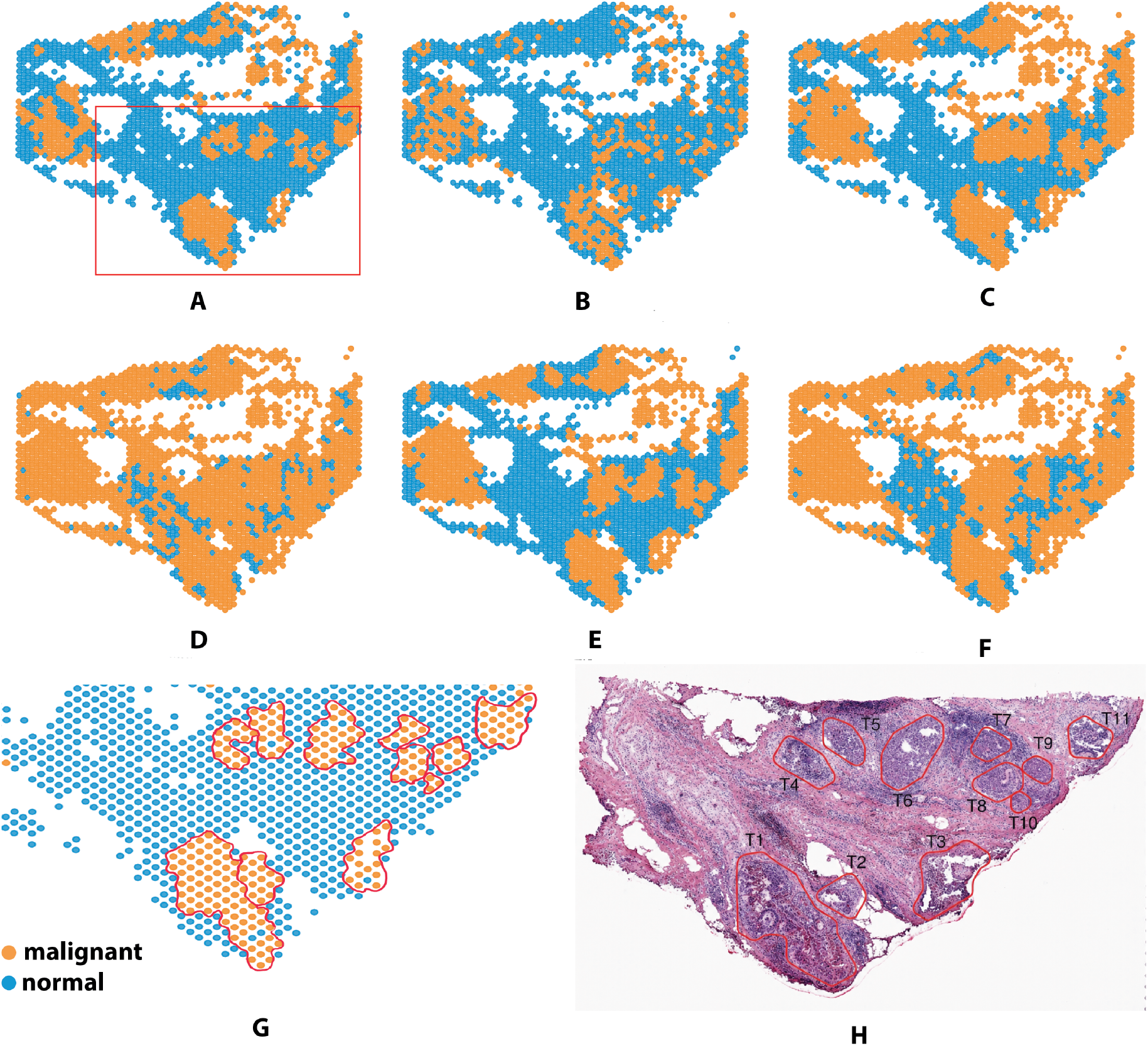
Spatial prediction comparison and histopathological validation on the DCIS2 Visium sample. (A–F) Tumor (orange) and normal (blue) regions predicted by (A) DeepMalignant, (B) DeepMalignant with constant edge weights, (C) CopyKAT, (D) scMalignantFinder, (E) Cancer-Finder, and (F) PreCanCell. (G) DeepMalignant prediction map annotated against the histopathological reference, showing that the predicted tumor regions recover the major annotated tumor foci. (H) Corresponding histopathology image with expert-annotated tumor regions, adapted from Wei et al. [20].

We then applied DeepMalignant to DCIS1, and found a similar pattern as DCIS2 (**SI Appendix Fig. S4**). As in DCIS2, DeepMalignant produced spatially coherent tumor regions, whereas the DeepMalignant with constant edge weight and all the four competing methods showed either fragmented malignant-spot regions, or overcalled malignant spots.

Together, these results demonstrated that DeepMalignant accurately distinguished malignant from normal spots in Visium-based spatial transcriptomic data, attributable to its joint integration of transcriptomic and copy number profiles. The spatially fragmented predictions observed in the constant-edge-weight variant, which lacked copy number profile input, underscored the indispensable complementary role of copy number variation signals alongside gene expression in achieving coherent and accurate malignant spot identification.

### Spatial cell-type identities and cell-cell communications in DCIS2 reveal contrasting fibroblast-to-epithelial signaling between normal and malignant epithelial cells

Since DeepMalignant provided us the capability of identifying the normal spots in 10x Visium data, we further integrated the normal spot prediction in DCIS2 with a matched scRNA-seq reference (**Methods: Cell-type annotation and spatial communication analysis in DCIS2**) to annotate each normal spot’s cell type compositions using RCTD [21]. **Figure 5A** showed the resulting pie chart for each spot, whereas each pie’s size was from the RCTD inferred weights for the corresponding cell type. The normal compartment was dominated by epithelial contributions across much of the tissue, whereas fibroblast-and myeloid-enriched mixtures were more prominent in localized regions, particularly in the central and lower-central portions of the section. Endothelial, B-cell, T-cell, and NK-cell contributions were comparatively sparse and spatially scattered. Tumor regions occupied spatially distinct areas of the section and were largely separated from these heterogeneous normal mixtures.

**Figure 5:**
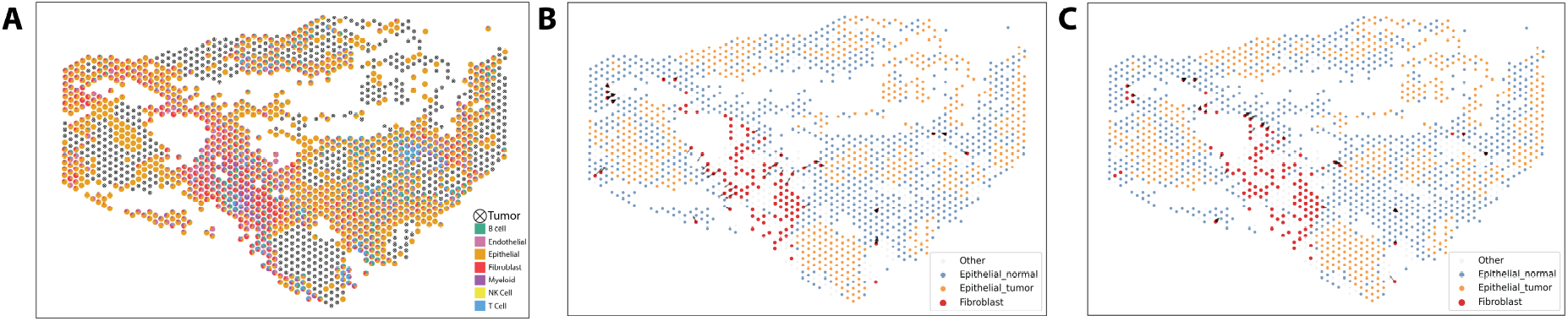
Spatial organization of the DCIS2 microenvironment and fibroblast-derived epithelial-directed communication. (A) RCTD-inferred cell-type composition of the normal spots predicted by DeepMalignant reveals spatial context in DCIS2. Normal spots are displayed as pie charts, where each slice represents the relative contribution of a reference cell type within that spot. Tumor spots are shown only for spatial context and are indicated by black crossed circles. Fibroblast-focused COMMOT spatial communication analysis for ligand–receptor axes C3–C3AR1 (B) and MIF–CD74/CD44 (C). Each point represents a spatial spot colored by cell type, with epithelial-normal spots shown in blue, epithelial-tumor spots in orange, fibroblasts in red, and other cell types in gray. Black arrows indicate fibroblast sender spots and their weighted epithelial-directed communication vectors, with arrow opacity and thickness reflecting communication strength.

We then further investigated cell-cell communication on DCIS2 by running COMMOT [22] given the annotation of the malignant and normal spots from DeepMalignant, which allowed us to look into the ligand-receptor pairs among different cell types (**Methods: Cell-type annotation and spatial communication analysis in DCIS2**). We found that fibroblast-derived C3 and MIF both directed signaling preferentially toward normal epithelial cells rather than tumor epithelial cells in the DCIS microenvironment, via the C3–C3AR1 and MIF–CD74/CD44 axes respectively (**Figure 5B-C**), reflecting a broader pattern in which DCIS tumor epithelial cells appear to disengage from homeostatic fibroblast paracrine circuits. For the C3–C3AR1 axis, we propose that the preferential signaling toward normal epithelial cells is driven by the loss of C3AR1 expression in tumor epithelial cells, a phenomenon previously attributed to epigenetic silencing through promoter methylation across multiple cancer types, while normal epithelial cells retain intact C3AR1 expression and remain the primary signal recipients [23].

For the MIF–CD74/CD44 axis, effective signal transduction requires cis-complex formation between CD74 and CD44 as an absolute prerequisite for downstream ERK1/2 activation [24, 25]; we speculated that in ER-positive/PR-negative DCIS2, CD74 was reduced in the tumor epithelial compartment, rendering the receptor complex non-functional and redirecting fibroblast MIF toward adjacent normal epithelium where CD74 expression was maintained [26, 27].

Together, these findings suggested that DCIS tumor epithelial cells might withdraw from fibroblast-derived homeostatic signaling through convergent receptor loss, pointing to stromal decoupling as a potentially early and coordinated feature of immune evasion in pre-invasive breast cancer.

### Ablation analysis highlighted contributions of copy number signal, attention, and gene selection

To quantify the contribution of individual components of DeepMalignant, we performed an ablation analysis in which key elements of the framework were removed or modified one at a time, including 1) replacing the edge weight as a constant number for all edges and thus remove copy number-based weighting; 2) using expression-only autoencoder trained on the same RNA feature set and thus remove graph-based neighbor aggregation and CNA weighting; 3) removing the most informative gene features (**Methods: Selection of highly correlated genes**). The comparison of the full model and each of these three settings is shown in **Table 1**.

The full model consistently achieved the best performance on both PDAC_A and PDAC_B, with F1 scores of 0.945 in both datasets. Recall reached 0.934 and 0.912, whereas precision reached 0.956 and 0.980 for PDAC_A and PDAC_B, respectively.

Removing copy number–based graph weighting resulted in a substantial reduction in performance, with F1 scores decreasing by 14% and 17% in PDAC_A and PDAC_B, respectively. Both precision and recall declined, indicating that copy number information contributes broadly to the discrimination between malignant and nonmalignant cells.

The largest performance degradation was observed when the attention mechanism was removed. In this setting, the F1 score decreased by 78% and 65% relative to the full model in PDAC_A and PDAC_B, respectively. In both samples, this degradation was accompanied by a marked loss of precision, indicating substantial overcalling of malignant cells when graph-based neighborhood aggregation was removed. These results suggested that, in the Moncada pancreatic cancer dataset, RNA features alone were insufficient to robustly separate malignant from non-malignant cells, whereas the graph attention framework provided essential contextual information that improved class separation in the latent space. This pattern was consistent with the lower precision observed in RNA-only approaches such as scMalignantFinder, PreCanCell, and Cancer-Finder across both samples. More broadly, these results indicated that, in datasets with substantial biological and technical noise, adaptive integration of neighborhood-derived information was critical for achieving stable and accurate predictions.

Finally, excluding the most informative gene features reduced the F1 score by 36% in PDAC_A and 16% in PDAC_B, demonstrating that these genes provided important signals for identifying malignant cells. This result also provided functional support for the attribution analysis, indicating that the genes identified through the gene–embedding correlation analysis were not merely associated with the learned representation, but also made an important contribution to its predictive utility.

Taken together, these ablation results demonstrated that multiple components contributed to the strong performance of DeepMalignant. Copy number–informed edge weighting improved the quality of the underlying graph representation, attention-based aggregation enhanced representation learning, and inclusion of highly informative gene features strengthened discriminative signals in the latent space. The pronounced performance degradation observed upon removal of the attention mechanism further indicated that graph-aware aggregation was particularly important in challenging datasets such as Moncada et al., where the biological signal was weaker and the data were noisier.

### Attribution analysis identified the most biologically relevant gene sets for breast cancer samples

On each sample of Gao et al., we further selected the top genes that were highly correlated with the GAT embeddings (**Methods: Selection of highly correlated genes**), and identified genes that appeared in the top lists of at least five embedding dimensions, forming a subset of recurrently important genes. We then conducted functional annotation and pathway enrichment analysis using the Database for Annotation, Visualization and Integrated Discovery (DAVID) [28] (**Dataset S2**) to further investigate whether these genes were indicative of the cancer type represented by each sample. The results supported the biological relevance of the recurrently important genes to breast cancer, with the most distinct signatures observed in DCIS1. Functional annotation of DCIS1-selected genes revealed two tightly coupled programs. The first was a hormone-responsive transcriptional program, evidenced by enrichment of the estrogen signaling pathway (HSP90AA1, TFF1, FKBP4, ESR1), response to estrogen (HSP90AA1, CCND1, ESR1), nuclear estrogen receptor binding (XBP1, ESR1), and endocrine resistance (CCND1, CDK4, ESR1). The second was a proliferative cell-cycle program, evidenced by enrichment of the cell cycle pathway (CCND1, CDK4, RAD21, YWHAZ, MAD2L1), cell division (CCND1, CDK4, RAD21, BIRC5, MAD2L1), G1/S transition of the mitotic cell cycle (CCND1, CDK4), and the spindle assembly check-point (BIRC5, MAD2L1). Both programs converged on the breast cancer and pathways in cancer terms (CCND1, CDK4, ESR1, HSP90AA1, BIRC5), confirming that the DCIS1-selected genes capture a canonical hormone-driven breast cancer program bridging endocrine signaling with accelerated cell cycle entry. TNBC1 was dominated by stress-response and chaperone-related genes such as HSP90AB1, HSPA1A/HSPA1B, and FKBP4. TNBC2 included genes such as CDKN2A, LGALS1, HDAC2, and CTSD, consistent with broader oncogenic and regulatory processes. TNBC3 showed enrichment for cell-cycle and oxidative stress programs, with representative genes including CDKN2A, MYBL2, NQO1, HMGA1, and YWHAZ. Overall, these results suggested that the genes highly correlated with the embedding capture biologically meaningful pathways relevant to breast cancer, with the clearest breast-specific enrichment observed in DCIS1 and broader malignant programs represented in the TNBC samples.

### Perturbation analysis revealed the contribution of CNA and transcriptional program to tumor-normal separation in multiple cancer types

To quantify the relative contributions of CNAs and transcriptional programs to the final tumor–normal separation in multiple cancer types, we performed a perturbation-based attribution analysis. In this analysis, we treated the final tumor–normal assignment obtained from the full model as the reference result, and then perturbed one component at a time while keeping all remaining model settings unchanged.

Specifically, we kept everything in our model the same except alternatively removing the following components: 1) CNA, genes related with 2) cell cycle, 3) DNA damage, 4) DNA repair, 5) invasion, 6) evading growth suppressors, 7) enabling replicative immorality, 8) genome instability and mutation, 9) resisting cell death, and 10) normal-enriched (**Methods: Perturbation-based contribution analysis**). Each component was assigned a score *S* which quantified the contribution from this component. A larger value of *S* indicated a larger contribution from the component. **Figure 6A-D** showed the perturbation scores across four datasets spanning breast, ovarian, and colorectal cancer.

**Figure 6:**
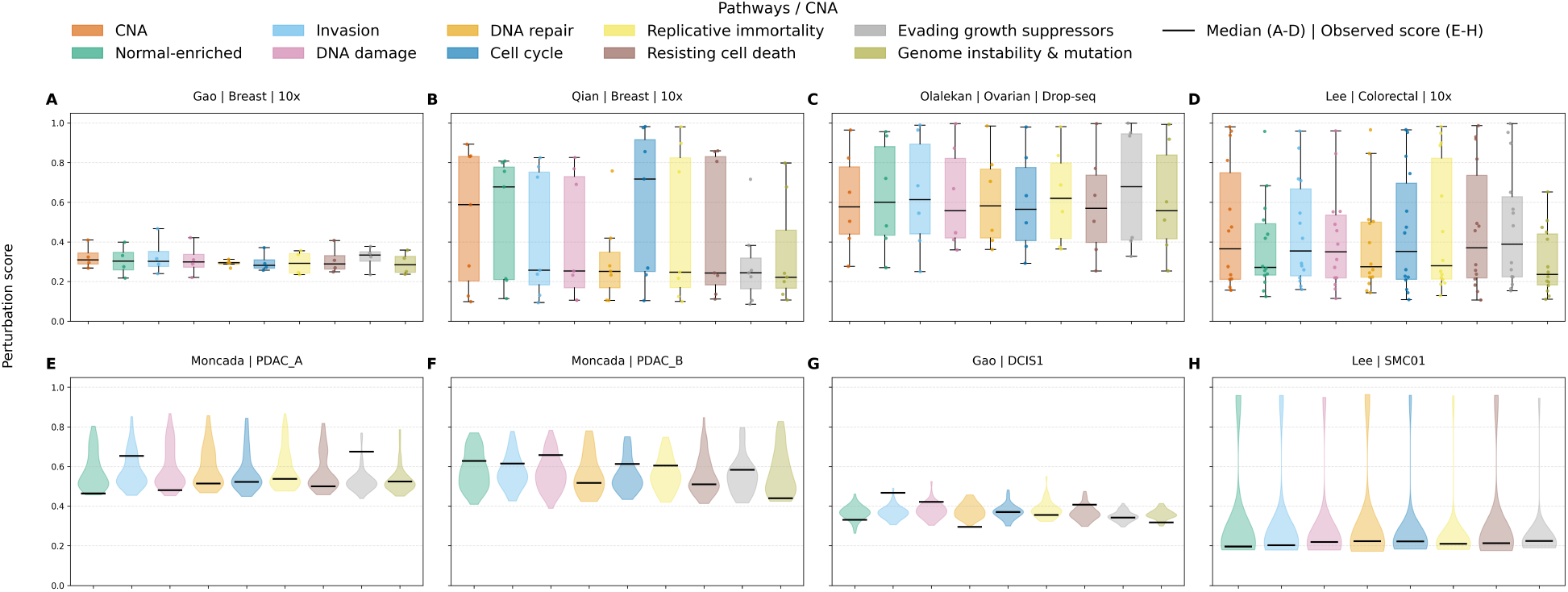
Perturbation and null-model analysis using V-measure on the final tumor–normal separation. Panels A–D summarize perturbation scores across datasets for (A) Gao et al. breast cancer, (B) Qian et al. breast cancer, (C) Olalekan et al. ovarian cancer, and (D) Lee et al. colorectal cancer. Panels E–H show null-model analyses for samples: (E) Moncada PDAC_A, (F) Moncada PDAC_B, (G) Gao DCIS1, and (H) Lee SMC01.

Among all four datasets, Qian et al. breast cancer dataset had the most contrastive perturbation scores among the ten components. In more detail, CNA, normal-enriched and cell cycle had much higher perturbation score than the rest of the components. This was expected and reflected the composition of the dataset, considering that Qian et al. dataset was a pan-cancer tumor microenvironment (TME) atlas focused on stromal cell heterogeneity. Specifically, CNA distinguished the rare malignant epithelial cells from the dominant stromal populations. Normal-enriched signatures captured the transcriptional similarity of the stromal majority to normal tissue. Cell cycle genes represented the most discriminative transcriptional feature of the malignant epithelial minority.

We then performed the null-model analysis for four selected cancer samples, Moncada et al. pancreatic cancer samples PDAC_A and PDAC_B, Gao et al. breast cancer sample DCIS1, and Lee et al. colorectal cancer sample SMC01 (**Figure 6E-H, Methods: Perturbation-based contribution analysis**). PDAC_A displayed significantly higher perturbation score for evading growth suppressors, whereas DCIS1 displayed a significantly higher perturbation score for invasion. The elevated perturbation score for evading growth suppressors in PDAC_A was consistent with the near-universal co-inactivation of CDKN2A/p16, TP53, and SMAD4 in pancreatic cancer [29–31], making this hallmark one of the strongest transcriptional discriminators between tumor and normal pancreatic cells. Conversely, the elevated invasion score in DCIS1 reflected the active transcription of ECM-remodeling and invasion-related gene programs in DCIS epithelial cells as they prime for stromal breach [32, 33], rendering invasion highly specific to the malignant population despite DCIS being a pre-invasive lesion.

## Conclusion

Accurate discrimination of malignant and normal cells is a prerequisite for reliable analysis of single-cell cancer transcriptomes, yet remains difficult across tumor types, experimental platforms, and data qualities. Here, we present DeepMalignant, an unsupervised multimodal graph representation learning framework that integrates RNA-derived malignant-cell signatures with CNA-informed cell–cell relationships to identify malignant cells from scRNA-seq data. By coupling transcriptional features with CNA-guided graph attention, DeepMalignant captures both dynamic malignant cell states and the more stable genomic structure shared across tumor cells.

Across 26 samples spanning breast, colorectal, pancreatic, and ovarian cancers generated on 10x Genomics, inDrop, and Drop-seq platforms, DeepMalignant consistently achieved a strong balance between precision and recall and showed greater cross-sample stability than existing RNA-based and CNA-based methods. Its advantage was especially clear in challenging settings, including pancreatic cancer and platform-shifted datasets, where expression-only methods tended to overcall malignancy and CNA-only methods showed substantial instability. Ablation analyses further demonstrated that both CNA-derived edge weighting and graph-based neighborhood aggregation are essential to performance, supporting the central design principle of multimodal graph integration.

The limited separation between tumor and normal cells observed in the inDrop pancreatic cancer dataset likely arises from a combination of platform-specific technical constraints and the intrinsic biological characteristics of pancreatic tumors. Compared to higher-sensitivity platforms such as 10x Chromium and Drop-seq, inDrop typically exhibits lower transcript capture efficiency, reduced gene detection per cell, and elevated dropout rates, which collectively diminish the signal-to-noise ratio and obscure subtle transcriptional differences. This limitation is particularly consequential in pancreatic cancer, where malignant ductal cells often closely resemble their normal counterparts at the gene expression level, and where the tumor microenvironment—comprising abundant stromal and immune cells—can further dilute tumor-specific signals. Additionally, the high heterogeneity of pancreatic tumors and the presence of low malignant cell fractions in some samples exacerbate the challenge of distinguishing tumor from normal cells using expression data alone. Nevertheless, DeepMalignant robustly and accurately distinguished the malignant and non-malignant cells in both PDAC_A and PDAC_B, thanks to its design of integrating both RNA-derived malignant-cell signatures with CNA-informed cell–cell relationships.

Beyond classification accuracy, DeepMalignant learned biologically meaningful representations. Attribution analyses showed that the latent space was associated with established cancer-related transcriptional programs, while application to matched spatial transcriptomics data recovered spatially coherent tumor regions that aligned closely with histopathological annotation. In addition, accurately separating the malignant and non-malignant spots in the spatial transcriptome data also enabled the cell-cell communication analysis contrasting the fibroblast-to-epithelial signaling between normal and malignant epithelial cells on a DCIS sample.

Together, these results establish DeepMalignant as a robust framework for malignant cell identification across diverse single-cell and spatial transcriptomic datasets. More broadly, they highlight the value of combining transcriptional and inferred genomic signals within graph-based representation learning to improve tumor annotation and to support down-stream studies of tumor heterogeneity, cell-cell communications, and microenvironmental organization.

## Methods

### Feature Extraction

Given a single cell RNA-seq dataset containing *N* cells, *G* genes and *D* genomic bins for copy number measurement, we construct a cell–cell graph integrating gene expression and copy number alteration (CNA) information.

#### RNA node features

Let *X*^raw^ ∈ *R*^*N*×*G*^ denote the raw UMI count matrix. We perform library size normalization followed by log transformation:eqn

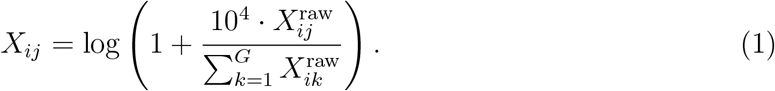

We then restrict the feature space to a curated set of cancer signature genes *S* derived from the scMalignantFinder framework [4] (the list of genes is in **Dataset S3**). These signatures capture gene expression programs characteristic of malignant cells and have been validated across multiple cancer datasets. The resulting node feature matrix iseqn

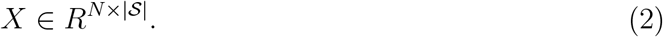

Each gene feature is standardized across cells using z-score normalization:

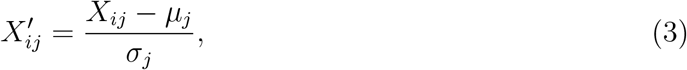

where *µ*_*j*_ and *σ*_*j*_ denote the mean and standard deviation of gene *j* across all cells.

#### CNA-based edge construction

Let *C* ∈ *R*^*N*×*D*^ denote the CNA matrix after filtering genomic bins with low variance:

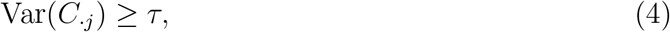

in which *τ* is a tunable variable whose default is 0.02 for eliminating genomic bins whose copy number does not vary among the cells. We construct a *k*-nearest neighbor (kNN) graph in CNA space using cosine similarity *d*_cos_(·,·):

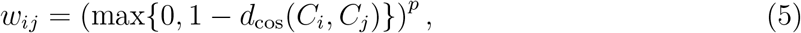

where *p* is a sharpening exponent. The resulting weighted adjacency matrix is denoted as *A* ∈ *R*^*N*×*N*^.

### Mathematical modeling and graph attention network

The overall framework is illustrated in **Figure 1**. RNA-derived features define node attributes *X* ∈ *R*^*N*×*d*^ (**Figure 1A**), while CNA similarity defines weighted edges *A* ∈ *R*^*N*×*N*^ (**Figure 1B**).

#### Graph attention encoder

We employ a two-layer graph attention encoder (**Figure 1C**). The first layer performs attention-based message passing:

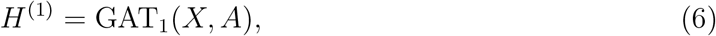

where the transformed node features are

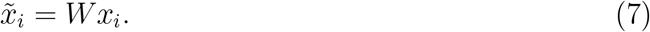

For each pair of connected nodes (*i, j*), attention scores are computed as

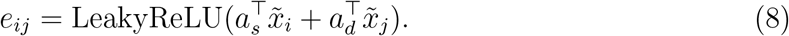

The CNA-derived edge weight *A*_*ij*_ directly modulates the attention score:

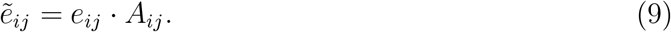

Row-wise softmax normalization produces attention coefficients:

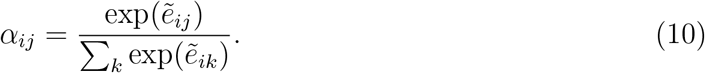

Node embeddings are then updated by aggregation:

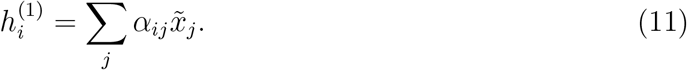

The second encoder layer applies a linear graph transformation without additional attention:

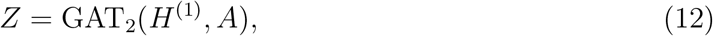

yielding latent embedding *Z* ∈ *R*^*N*×*r*^.

#### Decoder with weight tying

The decoder mirrors the encoder structure:

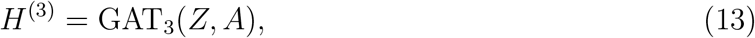

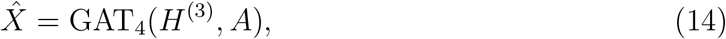

where decoder weights are tied to encoder weights such that

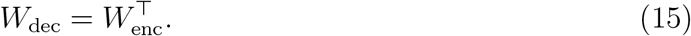

The decoder reuses the attention structure from the first encoder layer.

#### Loss function

The model is trained using a reconstruction loss combined with a graph-based contrastive loss:

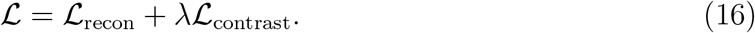

The reconstruction loss is defined as

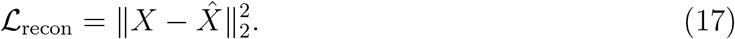

To prevent embedding collapse and enforce consistency among CNA-neighboring cells, we incorporate an InfoNCE contrastive loss:

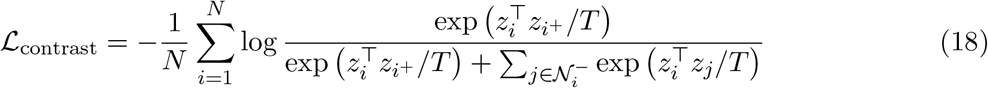

where *i*^+^ denotes a positive neighbor sampled from the K nearest neighbors (KNN) determined by cellular CNA cosine similarity, and 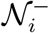 denotes a set of randomly sampled negative cells. Here, *T* is a temperature parameter controlling similarity scaling.

#### Model selection

During training, embeddings are clustered periodically using the Leiden algorithm (**Figure 1D**), and the model state yielding the highest silhouette score is selected for downstream malignant–normal classification.

#### Unsupervised malignant-cluster calling using CNA burden

To determine whether each cluster from Leiden algorithm belongs to the tumor or normal component, we investigate all cells’ CNA profiles inside each cluster. In more detail, let *C* ∈ *R*^*N*×*D*^ denote the CNA matrix whose cell order is corresponding to the clustered cell order by Leiden algorithm. We center CNA values per genomic segment by subtracting the segment-wise median across cells. For each cell *i*, we compute a CNA burden score

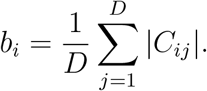

To stabilize the distribution, we apply a logarithmic transform:

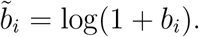

For each latent cluster *c*, we compute a cluster-level burden score

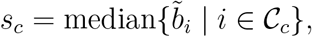

where *C*_*c*_ denotes the set of cells in cluster *c*.

We model the distribution of cluster-level scores {*s*_*c*_} using a two-component Gaussian mixture:

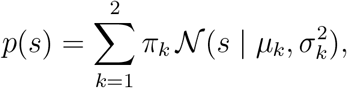

where *π*_*k*_ are mixture weights and *N* denotes the Gaussian density. Parameters are estimated via GaussianMixture implementation in scikit-learn.

The component with the larger mean *µ*_*k*_ is designated as the tumor component. In another word, suppose *µ*_1_ *< µ*_2_, then *µ*_T_ = *µ*_2_, *σ*_T_ = *σ*_2_, *π*_T_ = *π*_2_, and *µ*_N_ = *µ*_1_, *σ*_N_ = *σ*_1_, *π*_N_ = *π*_1_, in which *µ*_T_, *σ*_T_ and *π*_T_ represent the mean, standard deviation, and mixture weight for the tumor component, and *µ*_N_, *σ*_N_ and *π*_N_ represent the mean, standard deviation, and mixture weight for the normal component. For each cluster *c*, the posterior probability of its cells belonging to the tumor component is

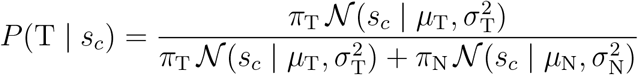

Cluster *c* is classified as tumor if

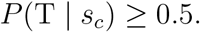

### Benchmark metrics

We evaluated tumor cell identification performance using precision, recall, and F 1 score, with the benchmark tumor–normal labels treated as ground truth. For each sample, cells predicted as malignant were considered positive predictions. True positives (TP) denote malignant cells correctly predicted as malignant, false positives (FP) denote normal cells incorrectly predicted as malignant, false negatives (FN) denote malignant cells incorrectly predicted as normal, and true negatives (TN) denote normal cells correctly predicted as normal.

Recall measures the fraction of true malignant cells recovered by a method and was defined as

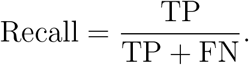

Precision measures the fraction of predicted malignant cells that were truly malignant and was defined as

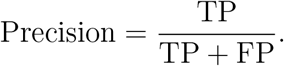

The F1 score summarizes the balance between precision and recall and was calculated as

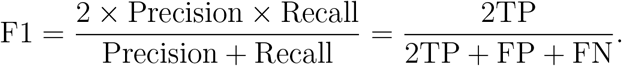

Metrics were calculated separately for each sample.

### Model parameter selection

See **SI Appendix, Research Design and Hyperparameter Choices**.

### Selection of highly correlated genes

To examine whether malignant and normal cells could be separated using the most informative transcriptomic features alone, we selected genes that were strongly associated with the DeepMalignant latent embedding. For each sample, we used the RNA feature matrix from the trained DeepMalignant input and the corresponding latent embedding matrix. For each gene and each latent embedding dimension, we computed the Pearson correlation between the gene expression values and the latent embedding values across all cells. This produced a gene-by-dimension correlation matrix. For each embedding dimension, genes were ranked by the absolute value of their Pearson correlation coefficient, and the top 20 genes were selected. The final highly correlated gene set was defined as the union of the top-ranked genes across all latent dimensions. Heatmap for Moncada et al. dataset is shown in (**SI Appendix Fig. S3**).

### Perturbation-based contribution analysis

We considered two classes of perturbations. Suppose the tumor–normal assignment obtained from the full model is *T*_0_, which will serve as the reference. First, to assess the contribution of CNA information, we replaced the CNA-derived edge weights with constant weights and recomputed the final tumor–normal assignment, yielding *T*_CNA removed_. The CNA contribution score was defined as

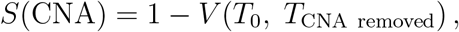

where *V* (·,·) denotes the V-measure between the perturbed and original final tumor–normal assignments. A larger value of *S*(CNA) (i.e., a smaller value of the V-measure) indicates that removing CNA-informed graph structure causes a larger change in the final tumor–normal separation, implying a stronger contribution of CNA-derived cell–cell relationships to the model output.

Second, to assess pathway-level transcriptional contributions, we perturbed one predefined pathway at a time. We used the cancer-related signature groups already incorporated into our feature design: *Cell Cycle, DNA damage, DNA repair, Invasion, Evading growth suppressors, Enabling replicative immortality, Genome instability and mutation, Resisting cell death*, and *Normal-enriched*. For a pathway *p*_*i*_, we removed all genes in that pathway from the RNA feature set, reran the model, and obtained the perturbed final tumor–normal assignment 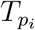. We then defined the pathway contribution score as

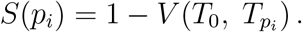

Thus, a larger value of *S*(*p*_*i*_) indicates that removing pathway *p*_*i*_ causes a larger change in the final tumor–normal separation and therefore suggests a stronger contribution of that pathway to the prediction.

To further evaluate whether the observed pathway perturbation effects were stronger than expected by chance, we constructed a matched-size null model for representative samples. For each pathway *p*_*i*_ containing |*p*_*i*_| genes, we repeatedly removed the same number of genes chosen uniformly at random from the original feature set, reran the full model, and recomputed the final tumor–normal assignment. For the *b*-th random perturbation, the null score was defined as

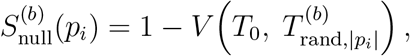

where 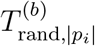 denotes the final tumor–normal assignment after the *b*-th random matched-size removal. This produced a null distribution of perturbation scores for each pathway. We then compared the observed pathway-specific perturbation score against its corresponding null distribution using the null mean and null standard deviation, which results in a z-score and an empirical p-value. This analysis controls for pathway size and allows us to assess whether the coordinated removal of genes within a biological pathway has a stronger effect than random removal of the same number of genes.

### Cell-type annotation and spatial communication analysis in DCIS2

For DCIS2, matched single-cell RNA-seq and Visium spatial transcriptomics data were used to annotate the normal tissue compartment and support downstream cell–cell communication analysis. The matched DCIS2 scRNA-seq count matrix was processed in Seurat [34] for quality control, normalization, dimensionality reduction, and clustering, and broad cell-type identities were assigned using SingleR [35]. These annotations defined major reference populations, including epithelial, T cell, B cell, myeloid, fibroblast, endothelial, and NK cell states. RCTD (version=2.2.1) was then applied to the Visium data using the annotated scRNA-seq profiles as the reference, and normal spots predicted by DeepMalignant were assigned cell-type compositions from the RCTD-inferred weights. For spot-level visualization and downstream annotation, the dominant cell type was defined as the reference class with the highest RCTD weight. Epithelial spots were further stratified into malignant and normal epithelial groups using DeepMalignant malignant–normal predictions.

### Cell–cell communication analysis

Cell–cell communication analysis was performed on the DCIS2 spatial transcriptomics dataset using COMMOT [22] (version=0.0.3). Raw 10x Visium data for the DCIS2 sample, aligned to the human reference genome (hg38) with Space Ranger, were loaded with Scanpy [36]. Spots outside the tissue were removed (`in_tissue` = 1), genes detected in fewer than 3 spots were discarded, and spots with fewer than 100 total counts were filtered out. Cell-type labels were then joined to the remaining spots by barcode, and spots lacking an annotation were excluded, yielding a set of annotated spots used as the cell-type grouping for all downstream cluster-level analyses.

Expression counts were normalized to a total of 10^4^ counts per spot and log-transformed (log(1+*x*)), with the unnormalized matrix retained. Ligand–receptor (LR) interactions were obtained from the human CellChat database [37] via COMMOT’s built-in interface, and the database was restricted to LR pairs whose ligand and receptor genes were each expressed in at least 5% of spots.

Spot-level signaling was computed with COMMOT’s `spatial_communication` function using the filtered CellChat database and a maximum spatial interaction distance threshold of 500 (in the units of the Visium spatial coordinates), with heteromeric complexes accounted for (`heteromeric = True`) and signaling aggregated at the pathway level (`pathway_sum = True`).

## Author contributions

X.M. and X.M.Z. designed research; P.B. performed research; P.B. and W.Y. analyzed data; all authors wrote the paper.

## Data and code availability

The single-cell RNA-seq datasets used for benchmarking DeepMalignant were obtained from the Curated Cancer Cell Atlas (3CA) [38], available at https://www.weizmann.ac.il/sites/3CA/. The same website also provided the ground truth malignant and normal cell labels derived from 3CA’s computational annotation pipeline, combining UMAP-based clustering, marker gene validation, and copy number aberration (CNA) inference via `infercna` [39]. The matched spatial transcriptomics data for the DCIS samples were obtained from the Gene Expression Omnibus (GEO, accession GSE181254) [20]. DeepMalignant is publicly available at https://github.com/compbio-mallory/DeepMalignant.

